## Supplementary figures for "Cryo-EM structure of *Helicobacter pylori* urease with a novel inhibitor in the active site at 2.0 Å resolution"

**Supplementary figures and legends**

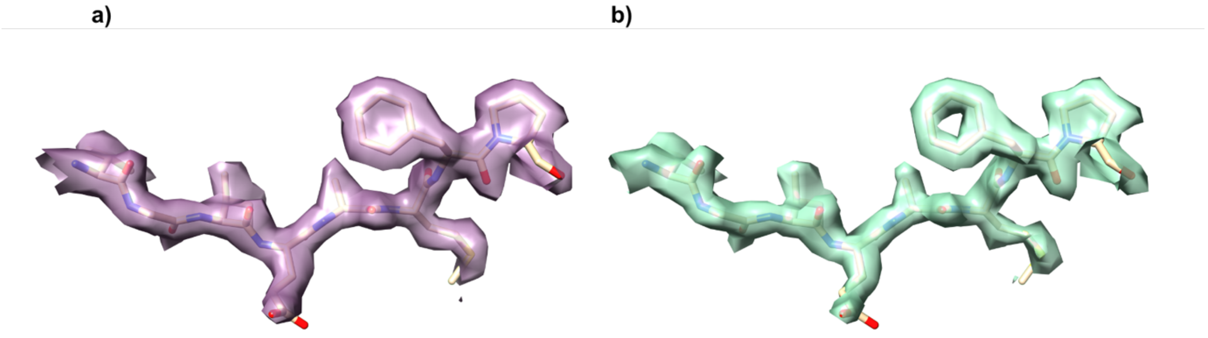


**Supplementary figure 1 – Density for U-SHA UreA residues A81-87 at 1σ contouring level. a)** Cryo-EM density at 2.09 Å resolution resulting from processing with the Relion package [22] (magenta). **b)** Density of the same region at 2.01 Å resolution after subsequent density modification with the resolve_cryo_em module from the Phenix package [23] (green). The model is depicted in sticks.

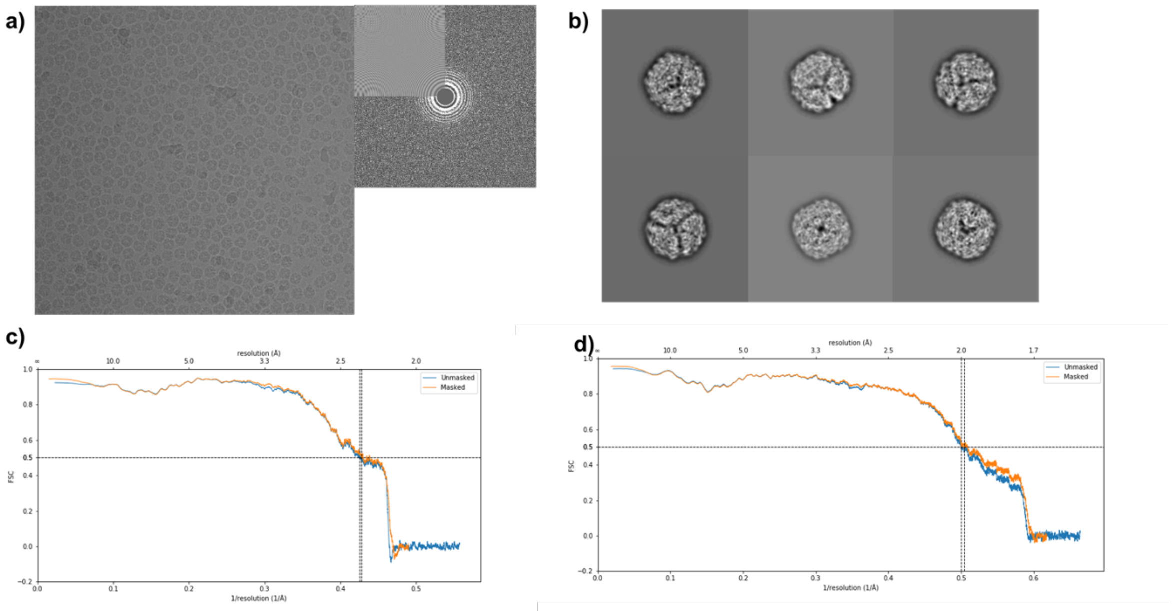


**Supplementary figure 2 – Cryo-EM data collection and processing. a)** Representative micrograph of U-BME particles suspended in vitreous ice at 1.077 Å/px and 2.8 µm estimated defocus. Only images exhibiting Thon rings beyond 5 Å were used for image processing. **b)** Representative reference-free 2D classes of U-BME exhibiting distinct features. **c)** Fourier Shell Correlation (FSC) curve for U-BME after density modification showing a nominal resolution estimation of 2.45 Å based on a model-to-map FSC with a cutoff of 0.5, which agrees well with the resolution reported using the map-to-map FSC with a cutoff of 0.143. **d)** Fourier Shell Correlation (FSC) curve for U-SHA after density modification showing a nominal resolution estimation of 2.01 Å based on a model-to-map FSC with a cutoff of 0.5, which agrees well with the resolution reported using the map-to-map FSC with a cutoff of 0.143.

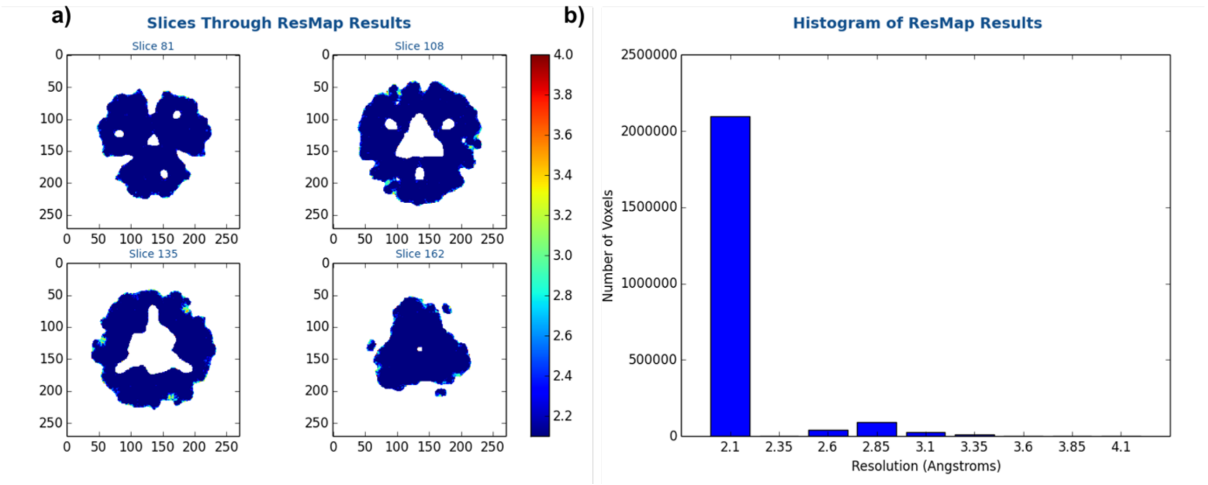


**Supplementary figure 3 – Local resolution estimates. a)** Slices depicting U-SHA resolution distribution as determined with the program Resmap [24] with dark blue areas showing higher resolution estimates and red areas showing lower resolution estimates. **b)** U-SHA voxel resolution distribution.

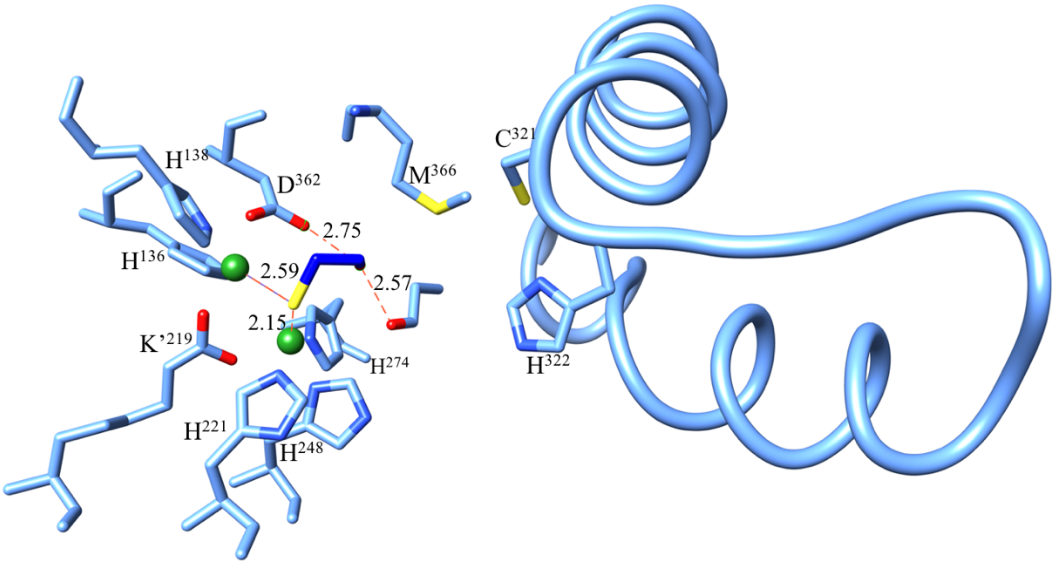


**Supplementary figure 4 – BME binding to *Helicobacter pylori* urease.** Urease flap region depicted in ribbon style with Cys321 and His322 on the right and active site residues in sticks on the right. Relevant distances in Å depicting the interactions of the BME molecule are as red dashed lines.

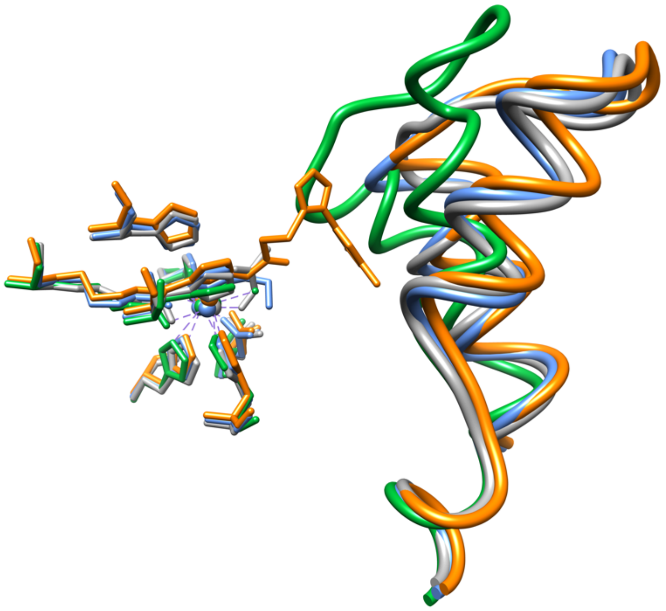

**Supplementary figure 5 – Comparison of all four *Helicobacter pylori* urease structures.** Urease flap regions depicted in ribbon (right) and active site residues in sticks (left). The four structures show high similarity for the active site, however they can be clustered according to differences in the conformation of the flap region with U-BME (blue, cryo-EM structure) and U-AHA (grey, crystal structure) clustered together and distinct conformations for the flap of U-SHA (more open, orange, cryo-EM structure) and U-APO (closed, green, crystal structure).
